# Knob architecture and hemoglobin composition shape recovery dynamics of *Plasmodium falciparum*–infected erythrocytes

**DOI:** 10.64898/2026.08.26.747262

**Authors:** Julian Czajor, Victor Lengyel, Cecilia P. Sanchez, Sabastian Dammrich, Anil K. Dasanna, Philipp R. Huppert, Leon Lettermann, Dmitry Fedosov, Fred Hamprecht, Ulrich S. Schwarz, Michael Lanzer, Motomu Tanaka

## Abstract

The deformability of the red blood cell (RBC) is essential for microcirculatory flow and is profoundly altered in hemoglobinopathies and during infection with *Plasmodium falciparum*. While many mechanical tests have been developed to probe RBC-mechanics, the dynamics of cell shape recovery following large deformations remains poorly characterized. Here, we integrate microfluidic constriction assays, ultrafast imaging, and computer simulations to quantify time-resolved shape recovery of individual erythrocytes. We show that parasite infection is the primary determinant of RBC viscoelastic behavior. In wild-type (HbAA) erythrocytes, the relaxation time increases progressively from ring to trophozoite to schizont stages, consistent with parasite-induced membrane stiffening and enhanced membrane– cytoskeleton coupling. In contrast, sickle trait (HbAS) erythrocytes exhibit a distinct response: although deformation becomes increasingly irreversible during parasite maturation, the relaxation time after constriction remains largely unchanged. Analysis of a mutant parasite line with enlarged and sparsely distributed knobs revealed a significant increase in relaxation time, demonstrating that knob architecture modulates recovery kinetics. Together, these findings suggest that the coupling between membrane and cytoskeleton, which is strongly changed by the establishment of the knobs during an infection with *Plasmodium falciparum*, should have a strong detrimental effect on microcirculatory flow, which is however weakened by the sickle cell trait.

## Introduction

Red blood cells are central to human physiology, acting as carriers of respiratory gases (Lux, 2016, Mohandas and Gallagher, 2008). They take up oxygen in the lungs and deliver it to peripheral tissues, where it is exchanged for carbon dioxide. The cells then return to the lungs to release carbon dioxide and reload with oxygen, sustaining the continuous cycle of gas transport. In the course of circulation, red blood cells, which measure about 7–8 µm in diameter, must repeatedly traverse narrow constrictions, such as microcapillaries (diameter 3.0–4.0 µm) and splenic interendothelial slits (which are as narrow as 0.3–1.0 µm). Passage through these geometrical constraints requires erythrocytes to undergo large, reversible deformations and to rapidly recover their shape upon exit.

The extraordinary capability to undergo such large deformations arises from a combination of geometric, cytoplasmic, and membrane mechanical properties. Erythrocytes exhibit a biconcave discoid shape with a high surface-to-volume ratio, providing excess membrane area for bending without significant membrane stretching (Lim H. W. et al., 2002). Their cytoplasm is a concentrated hemoglobin solution that behaves as a Newtonian fluid with a viscosity about five times as large as the one of the surrounding plasma (Fedosov et al., 2014). Most critically, the cell envelope is a composite structure consisting of a fluid lipid bilayer coupled to an elastic membrane skeleton, which endows the cell with high resistance to shear but low resistance to bending. The membrane skeleton is composed of α- and β-spectrin heterodimers assembled head-to-head into tetramers, which are interconnected at their N-termini by short actin protofilaments forming around 35,000 junctional complexes, and linked to the lipid bilayer at their C-termini via ankyrin–band 3 (anion exchanger 1) complexes and protein 4.1–glycophorin C interactions (Lux, 2016). This quasi-hexagonal network with a typical distance of 70 nm between the nodes behaves as an entropic spring lattice, defining the in-plane shear modulus, while the lipid bilayer largely determines the bending modulus. Importantly, this mechanical system also depends on ATP-concentration and it has been shown that the membrane fluctuations below 10 Hz frequency are active in nature, i.e. mainly driven by ion channels, while the membrane fluctuations above 10 Hz frequency correspond to passive thermal motion (Turlier and Betz, 2019, Turlier et al., 2016). Together, membrane dynamics, the coupling between membrane and cytoskeleton, and the hemoglobin solution determine the dynamical properties of RBCs. A recent study with optical tweezers on RBC-mechanics in fact identified three relaxation times after stretching, with a fast time scale of 0.01-0.1s, an intermediate time scale of 4s, and a slow time scale of 70s, which might correspond to these three elements (Gironella-Torrent et al., 2024).

During aging, erythrocytes progressively lose their ability to deform and are ultimately removed from circulation, primarily in the spleen, where mechanical filtration through interendothelial slits imposes stringent deformability thresholds. Loss of deformability also characterizes erythrocytes carrying hemoglobinopathies such as the sickle cell trait (Hebbel, 1991). The sickle cell trait arises from a valine-for-glutamic-acid substitution at position 6 of the β-globin chain and occurs in individuals who inherit one mutated (HbS) and one wild type (HbA) β-globin gene. Consequently, HbAS erythrocytes contain a mixture of HbA and HbS, which assemble with α-globin chains into heterotetrameric hemoglobin. As a result of this single amino acid substitution, sickle trait erythrocytes (HbAS) exhibit a measurable stiffening of their membrane, reflected in increased effective membrane tension, bending modulus, membrane confinement, and cytosolic viscosity (Fröhlich et al., 2019, Hebbel, 1991). Comparable, yet mechanistically distinct, biomechanical aberrations also occur in erythrocytes infected with the human malaria parasite *Plasmodium falciparum* (Fröhlich et al., 2019). Transmitted by the bite of an infected Anopheles mosquito, the parasite initially infects hepatocytes before entering the bloodstream and invading erythrocytes, where it establishes a ∼48-hour intraerythrocytic replication cycle. During this cycle, parasites develop from ring stages (0-20 hours post invasion (hpi)) to trophozoites (20-36 hpi) and schizonts (36-40 hpi), culminating in the release of invasive merozoites that perpetuate infection. Because the sickle cell trait protects carriers from the effects of a malaria infection, the two conditions must be closely linked.

In both hemoglobinopathic and infected erythrocytes, membrane stiffening has been strongly associated with elevated oxidative stress, arising either from HbS instability and reactive oxygen species generation or from parasite metabolism and hemoglobin digestion (Cyrklaff et al., 2011, Cyrklaff et al., 2016, Gwozdzinski et al., 2026). Oxidative modifications impact membrane skeletal architecture and bilayer–cytoskeleton coupling (Gwozdzinski et al., 2026). For example, oxidative stress promotes band 3 phosphorylation, clustering, and partial detachment from ankyrin, thereby weakening its vertical interactions with spectrin and leading to partial uncoupling of the cytoskeleton from the lipid bilayer (Ferru et al., 2011, Arashiki et al., 2013). Concurrently, spectrin undergoes oxidative crosslinking and reduced conformational flexibility, increasing the effective shear modulus of the membrane skeleton. In infected erythrocytes, oxidative stress additionally perturbs parasite-driven remodeling processes. For example, oxidative stress interferes with actin reorganization and protein export (Cyrklaff et al., 2011, Cyrklaff et al., 2016, Kilian et al., 2015). During intraerythrocytic development, *P. falciparum* exports a large repertoire of proteins that extensively reconfigure the host cell membrane and cytoskeleton (Jonsdottir et al., 2021, Maier et al., 2009). During early ring stage the ring-infected erythrocyte surface antigen (RESA) is released, phosphorylated and binds to β-spectrin, stabilizing the spectrin tetramer (Mills et al., 2007, Pei et al., 2007). Furthermore, actin is mined from junctional complexes and reorganized into elongated, dynamic filaments that facilitate vesicular trafficking of parasite-derived proteins across the host cytoplasm (Cyrklaff et al., 2011, Cyrklaff et al., 2016). This actin mining reduces junctional connectivity and alters the topology of the spectrin network, thereby weakening its intrinsic elasticity and modifying its strain response.

Concomitantly, parasite-encoded proteins such as the knob associated histidine-rich protein (KAHRP) and adhesins (such as P. falciparum erythrocyte membrane protein 1 (PfEMP1)) assemble into multiprotein complexes at the cytoplasmic face of the membrane, forming electron-dense protrusions known as knobs (Wiser, 2023). Knobs serve as platforms for the display of parasite adhesins, enabling infected erythrocytes to adhere to endothelial receptors and sequester within the microvasculature, thereby avoiding splenic clearance (Wiser, 2023). Knobs mechanically couple to spectrin—predominantly at junctional nodes (Sanchez et al., 2021)—and introduce localized stiffening and strain hardening of the membrane (Fröhlich et al., 2019). The resulting composite membrane exhibits increased shear modulus and reduced deformability, while simultaneously providing anchoring platforms for cytoadhesive interactions with endothelial receptors under flow. This remodeling is spatially heterogeneous, leading to anisotropic mechanical behavior and altered membrane fluctuation spectra.

In parallel, the growing parasite occupies an increasing fraction of the intracellular volume and consumes hemoglobin, thereby modifying cytosolic rheology (Waldecker et al., 2017, Lew et al., 2003). The combined effects of increased internal viscosity, reduced excess membrane area due to progressive cell rounding, and reinforced membrane–cytoskeleton coupling contribute to the marked loss of deformability observed in late-stage infected erythrocytes. Together, these processes fundamentally alter the mechanical phenotype of the cell, impairing its ability to traverse narrow capillaries and splenic slits while promoting sequestration within the microvasculature.

Given the critical role erythrocyte deformability plays in human health and disease, a broad spectrum of techniques—including ektacytometry, micropipette aspiration, microfluidics, atomic force microscopy, microfiltration, optical tweezers, imaging flow cytometry, and computational modeling—has been employed to quantify mechanical properties and elucidate underlying molecular mechanisms (Depond et al., 2019). These approaches have established key parameters such as membrane shear modulus, bending rigidity, effective membrane tension, and cytosolic viscosity, and have provided important insights into how these quantities are altered in hemoglobinopathies and during *P. falciparum* infection. These quantitative insights also form the basis of computational approaches to better understand the mechanics of malaria-infected RBCs (Dasanna et al., 2020). However, most of these techniques probe erythrocytes either under quasi-static conditions or in simplified mechanical configurations that do not fully recapitulate the complex, time-dependent deformations experienced in the microcirculation or the changes occuring in disease conditions like hemoglobinopathies or malaria infections. As a result, critical dynamic properties—such as relaxation timescales, hysteresis during repeated deformation cycles, and the extent of irreversible mechanical remodeling—remain insufficiently characterized.

In order to address physiologically relevant constriction geometries for both red and white blood cells, microfluidic setups have been developed that can recapitulate the relevant spatial and temporal dimensions (Peng et al., 2026). For example, the role of ATP-concentration for wildtype RBC relaxation dynamics has been demonstrated with a catch-load-launch device (Ito et al., 2017). To investigate the transit times in the spleen during malaria infection, a microfluidic setup has been combined with computer simulations (Li et al., 2024, Li et al., 2018, Moreau et al., 2023, Pivkin et al., 2016). Image data from passage through a microfluidic constriction has been recently used to train a neural network to detect malaria infections (Rademaker et al., 2023). However, a quantitative analysis of the relaxation dynamics after exit from a constriction and a quantitative framework linking transient cell shapes to underlying mechanical parameters is still lacking, particularly for the pathophysiological conditions of HbAS, malaria infection and their combination. Here, we address this gap by combining microfluidic constriction assays with ultra-fast imaging and computer simulations to directly resolve the full deformation–recovery cycle of individual erythrocytes. This approach enables us to extract not only steady-state measures of deformability, but also dynamic observables, including entry and transit times, relaxation kinetics, and shape memory effects. This allows us to identify distinct biomechanical signatures of uninfected and infected wild type (HbAA) and HbAS erythrocytes, and to relate these signatures to specific molecular alterations of the membrane skeleton and parasite-induced remodeling.

## Materials and Methods

### P. falciparum culture

The *P. falciparum* line FCR3 (also referred to as IT4) and SR1 were maintained in continuous culture. The blood-stage cultures containing HbAA or HbAS erythrocytes were maintained at a hematocrit of 3.5% and a parasitemia below 5% in RPMI 1640 medium supplemented with 25 mM HEPES, 2 mM L-glutamine, 100 µM hypoxanthine, 20 µg mL^-1^ gentamicin, and 10% (v/v) heat-inactivated human AB+ serum. Cultures were incubated at 37°C in a gas mixture of 3% CO_2_, 5% O_2_, and 92% N_2_, with relative humidity maintained at 96%. Parasite cultures were synchronized using a combination of sorbitol and heparin (Kobayashi and Kato, 2016). Briefly, sorbitol treatment was first applied to achieve partial synchronization by selectively lysing mature parasite stages. Subsequently, heparin (30 U ml^-1^) was added to inhibit merozoite invasion. Once the majority of parasites reached the schizont stage, heparin was removed to allow synchronized invasion. After 4 h, heparin was reintroduced to prevent further invasion. Repetition of this procedure over multiple cycles resulted in highly synchronized cultures within a narrow time window of approximately 4 h. For microfluidic experiments, tightly synchronized cultures were prepared with parasitemia levels of up to 15%.

### Microfluidics and ultrafast imaging

The microfluidic chambers were fabricated in-house using polydimethylsiloxane (PDMS). Each device contained 15 parallel channels, each with a total length of 190 µm. PDMS was cast onto a SU-8 mold, cured at 120°C for more than 2 h, and subsequently left to rest overnight at room temperature. The cured PDMS slab and a glass substrate were then treated with O_2_ plasma for 30 s and irreversibly bonded. Erythrocyte suspensions were introduced into the device at a flow rate of 0.5 µL min^−1^ using a neMESYS low-pressure module (CETONI GmbH, Korbußen, Germany). Cells entering the device were guided into the constriction zone, which had dimensions of 3 µm × 4 µm × 20 µm (width × height × length). Upon exiting the constriction, cells entered a relaxation zone measuring 10 µm × 4 µm × 165 µm, where shape recovery was monitored. The microfluidic device was mounted on an Axio Observer Z.1 inverted microscope (Zeiss, Oberkochen, Germany) equipped with a FastCam Mini AX50 high-speed camera (VKT Video Kommunikation GmbH, Pfullingen, Germany). Images were recorded from a region of interest (724 × 64 pixels) at a frame rate of 36,000 frames per second (corresponding to 27 µs per frame), as the rapid shape recovery dynamics could not be resolved at conventional frame rates (e.g., 33 fps).

### Image analysis

Each recorded movie contained up to 10,000 frames, with pixel intensities stored in grayscale (0–255). To improve image quality, noise was reduced and intensity variations were corrected by applying denoising, leveling, and normalization procedures (Chambolle, 2005). These preprocessing steps also facilitated the removal of static features, such as PDMS channel walls. Cell shapes were extracted by binarization using the Ilastik software and its integrated pixel classification framework (Berg et al., 2019). For training, up to 30 labels were manually assigned per frame, and approximately every 100th frame was used to train the classifier. In total, 333 movies were included in the training dataset to enable robust automated segmentation across varying imaging conditions. Based on the binarized images, the projected two-dimensional shape of each cell was approximated by fitting an ellipse using the OpenCV library (Fitzgibbon et al., 1999). This approach provided quantitative descriptors of cell geometry for subsequent analysis.

### Computer simulations

Fluid flow is modelled by the smoothed dissipative particle dynamics (SDPD) method, which is a Lagrangian discretization of the Navier-Stokes equations (Español and Revenga, 2003, Müller et al., 2015). The SDPD fluid is composed of N particles that interact via pairwise conservative, dissipative (both translational and rotational) and random forces. For more details about the SDPD algorithm, we refer the reader to (Müller et al., 2015, Dasanna et al., 2021).

RBC is modelled as a two-dimensional triangular meshwork of N_v_ vertices, *N*_e_ edges and *N*_t_ triangles (Fedosov et al., 2010). The total potential energy is:

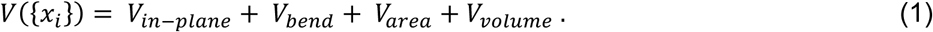

The first term is the in-plane elastic energy of the spring network:

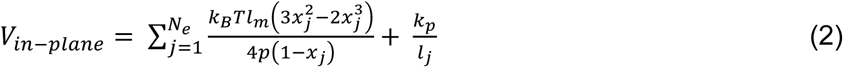

Both terms together act as a spring-like potential. In the first part, *p* is the persistence length, *l*_m_ is the maximum extension of the spring, and *x_jj_* = *l_jj_*/*l_m_*. In the second part, *k_p_* is the repulsive coefficient.

The second term in Eq. (1) represents the bending energy:

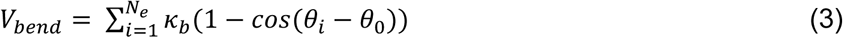

where *κ*_b_, *θ*_i_ and *θ*_0_ are the bending coefficient, the angle between two neighboring faces that share the same edge and the preferred angle, respectively.

The last two terms in Eq. (1) correspond to surface area and volume constraints:

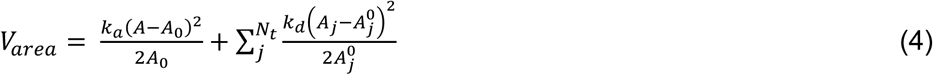

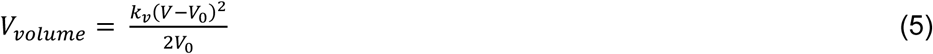

where *k_a_* and *k_d_* are the global area and local area constrain coefficients, and *k_v_* is the volume constraint coefficient. 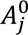 is the desired area of triangle j, and *A*_0_ and *V*_0_ are the desired area and volume of the RBC membrane.

The size of RBC is defined as 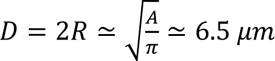. SDPD fluid particle density is set to be 6. RBC bending coefficient is set to 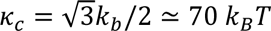 and shear modulus to *μ*_0_≃ 5*μN*/*m*. Channel walls are modelled by frozen SDPD particles. We distinguish the fluid inside and outside the RBC. Appropriate SDPD interactions between the fluid and RBC or wall particles are chosen to impose no-slip boundary conditions at the RBC surface and the walls.

### Preparation of Exposed Membranes and STED Microscopy

*P. falciparum*-infected erythrocytes (SR1 strain) were immobilized on functionalized glass-bottom culture dishes (MatTek Corporation) as previously described (Sanchez et al., 2021). Exposed membranes were incubated overnight at 4°C with rabbit anti-KAHRP antibody, followed by incubation with an Abberior STAR RED-conjugated anti-rabbit secondary antibody for 40 min at room temperature. All antibody incubations and washing steps were performed in PBS containing 3% bovine serum albumin (BSA).

Super-resolution images were acquired using a STED/RESOLFT microscope (Abberior Instruments, Germany) equipped with 488, 594, and 640 nm excitation lasers and a 775 nm STED depletion laser, mounted on an Olympus microscope with a 100× oil-immersion objective (UPLSAPO, NA 1.4, WD 0.13 mm). Fluorescence emitted by the Abberior STAR RED-conjugated secondary antibody was detected using the 640 nm excitation channel. Images were acquired with a pixel size of 15 nm and a pixel dwell time of 10 µs. Two-dimensional STED images were deconvolved using Imspector software (Abberior Instruments GmbH) with the Richardson–Lucy algorithm, applying the default settings and a regularization parameter of 1 × 10⁻¹⁰.

### Western Blot Analysis

Western blot analysis of SR1 cell extracts was performed as previously described (2). Briefly, trophozoite-stage parasites were enriched by magnetic purification and lysed in 10 mM phosphate buffer supplemented with protease inhibitors (Halt™ Protease Inhibitor Cocktail, Thermo Scientific). Membrane fractions were collected by centrifugation and resuspended in protein loading buffer (250 mM Tris-HCl, pH 6.8, 3% SDS, 20% glycerol, and 0.1% bromophenol blue), followed by brief sonication. The lysate was centrifuged at 17,000 × *g* for 30 min at 4°C, and the resulting supernatant was analyzed by SDS-PAGE.

Proteins were transferred onto a polyvinylidene difluoride (PVDF) membrane using an iBlot™ 2 transfer system. Membranes were probed with the following primary antibodies: mouse anti-KAHRP (1:1,000 dilution) and rabbit anti-PFE1605w (1:1,000 dilution; gift from H.-P. Beck). Goat anti-rabbit horseradish peroxidase (HRP)-conjugated antibody (1:10,000 dilution; Abcam) was used as the secondary antibody. All antibodies were diluted in PBS containing 1% (w/v) BSA. Immunoreactive signals were detected using a C-DiGit blot scanner (LI-COR Biosciences).

## Results

### Microfluid chamber design and analytical pipeline

Figure 1 depicts the microfluidic chamber, consisting of 15 constrictions arranged in parallel, each measuring 20 µm in length with a cross section of 4 µm × 3 µm. The details of integration of the self-developed microfluidic chip and the ultrafast imaging system are presented in Figure S1. To quantitatively analyze erythrocyte deformation and recovery within this microfluidic system, we established a machine learning–based image analysis pipeline utilizing Ilastik (Figure 2) (Berg et al., 2019). Raw images were manually annotated to label cells and background (in yellow and blue, respectively, in Figure 2a) providing training data for supervised machine learning–based segmentation (see Figure S2 for more detail). This approach enabled robust binarization and reliable extraction of cell contours across varying contrast conditions.

**Figure 1.**
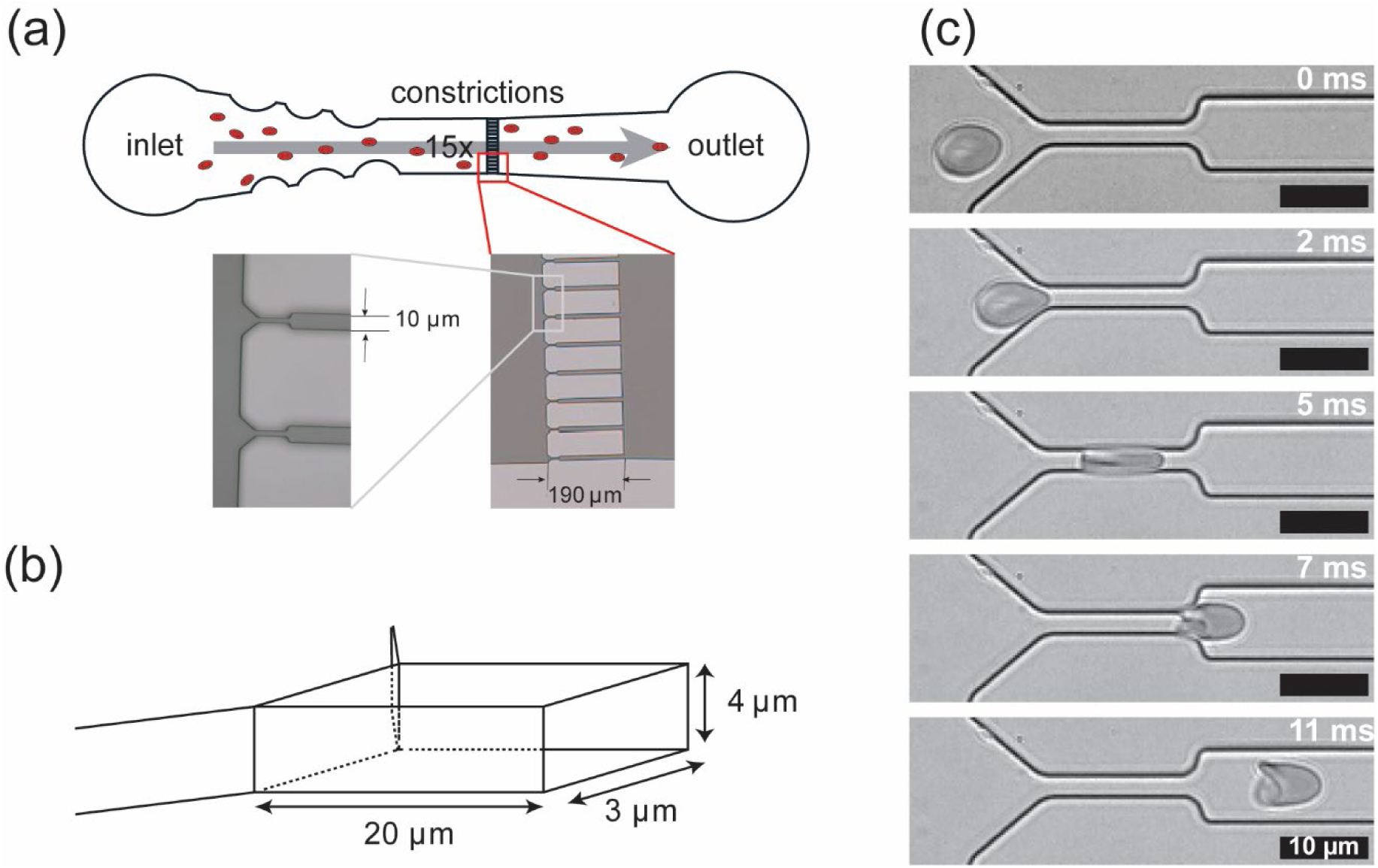
Microfluidic channels for ultrafast imaging of shape recovery of healthy and infected erythrocytes. (a) Coarse structure of microfluidic channels designed for this study. (b) Detailed structure of a constriction channel with a cross-section of 3 µm x 4 µm. (c) Typical snapshot images of uninfected erythrocyte passing through constriction. Ultrafast imaging (27 µs per frame) allows real-time imaging of the entry and passage of individual cells through the constriction zone mimicking dimensions and flow velocity of microvasculature, which is followed by viscoelastic shape recovery. Released cell was allowed to relax in 165 µm-long relaxation channels.

**Figure 2.**
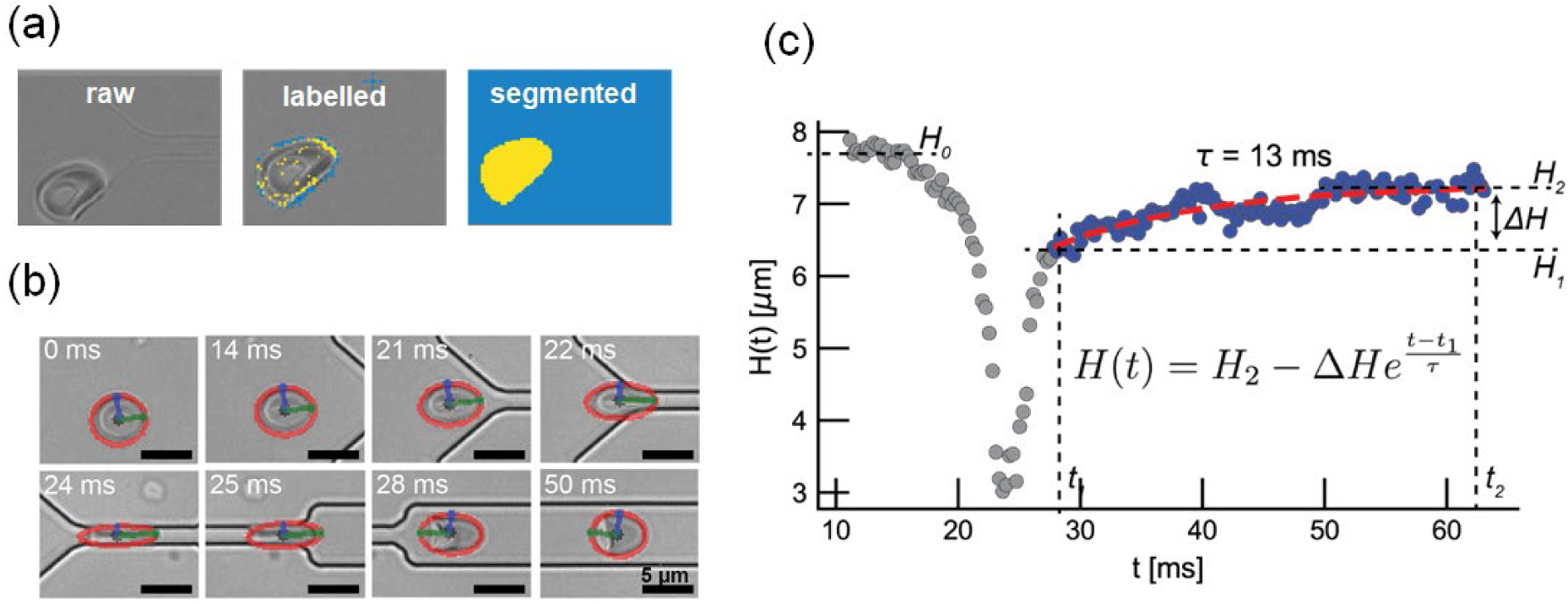
Flow of the analysis of experimental data. (a) Cell shape was extracted by the machine learning-assisted analytical routine. (b) 2D-projected cell shape was approximated as an ellipse (red), whose major radius and minor radius were labeled in green and blue, respectively. Cells exhibited significant rotations were excluded once the angle between the flow direction and the major axis becomes larger than 50 ° (Figure S2 for more detail). (c) Width *H* of an uninfected erythrocyte plotted as a function of time *t*. *H*_0_; initial, non-perturbed width, *H*_1_; width at the time point of complete exit, and *H*_2_; final width after traveling approx. 135 µm from exit. The viscoelastic shape recovery was fitted with an exponential function, yielding relaxation time *τ*.

To parameterize temporal changes in cell shape, ellipse fits were applied to segmented cell contours, yielding the major and minor axes (indicated in green and blue, respectively, in Figure 2b). Although not all cells recovered a round or discoid shape upon exiting the constriction—some exhibited a parachute-like phenotype (Figure 2c)—an ellipse-based description was adopted for quantitative analysis. This simplification is justified by the marked reduction in parachute-like morphologies upon infection with *P. falciparum*, with their frequency decreasing to below 20% at the trophozoite stage and to less than 5% at the schizont stage (Figure S3). Since our primary objective was to resolve differences in shape recovery dynamics during parasite maturation and associated membrane remodeling, the ellipse approximation provides a consistent and tractable metric across all conditions.

The minor radius was used to monitor the temporal change in cell width, *H*(*t*) = 2 × (minor radius), from which three characteristic widths were extracted at defined time points: *H*_0_, the initial unperturbed width prior to entry; *H*_1_, the width at the time point of complete exit; and *H*_2_, the final width measured approximately 135 µm downstream of the exit, corresponding to ∼80 % of the relaxation zone. Cells exhibiting significant rotational motion – typically arising from non-specific pinning at the channel exit - were excluded when the angle between the flow direction and the major axis exceeded 50° (Figure S2).

To quantify recovery kinetics, we defined a characteristic relaxation time, *τ*, obtained by fitting a simple exponential function to the time-dependent cell width *H(t)*, starting from *t*_1,_ the time point of complete exit:

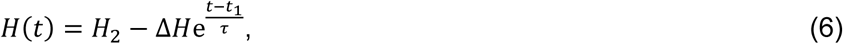

where Δ*H* = *H*_2_ − *H*_1_. For example, fitting the data shown in Figure 2c (dashed line) yields a relaxation time of *τ* = 13 ms.

### Differential shape recovery dynamics between uninfected HbAA and HbAS erythrocytes

As proof-of-principle for our experimental set-up, we investigated the deformability and shape recovery of sickle cell trait HbAS versus wild-type HbAA erythrocytes. As HbAS erythrocytes are known to be mechanically stiffer than HbAA cells, as demonstrated in multiple independent studies (Zheng et al., 2015, Maciaszek and Lykotrafitis, 2011, Fröhlich et al., 2019), we hypothesized that they exhibit altered recovery dynamics in our assay. Both cell populations predominantly displayed the characteristic discocyte morphology, indicating no gross morphological abnormalities under the experimental conditions (Figure 3a).

**Figure 3.**
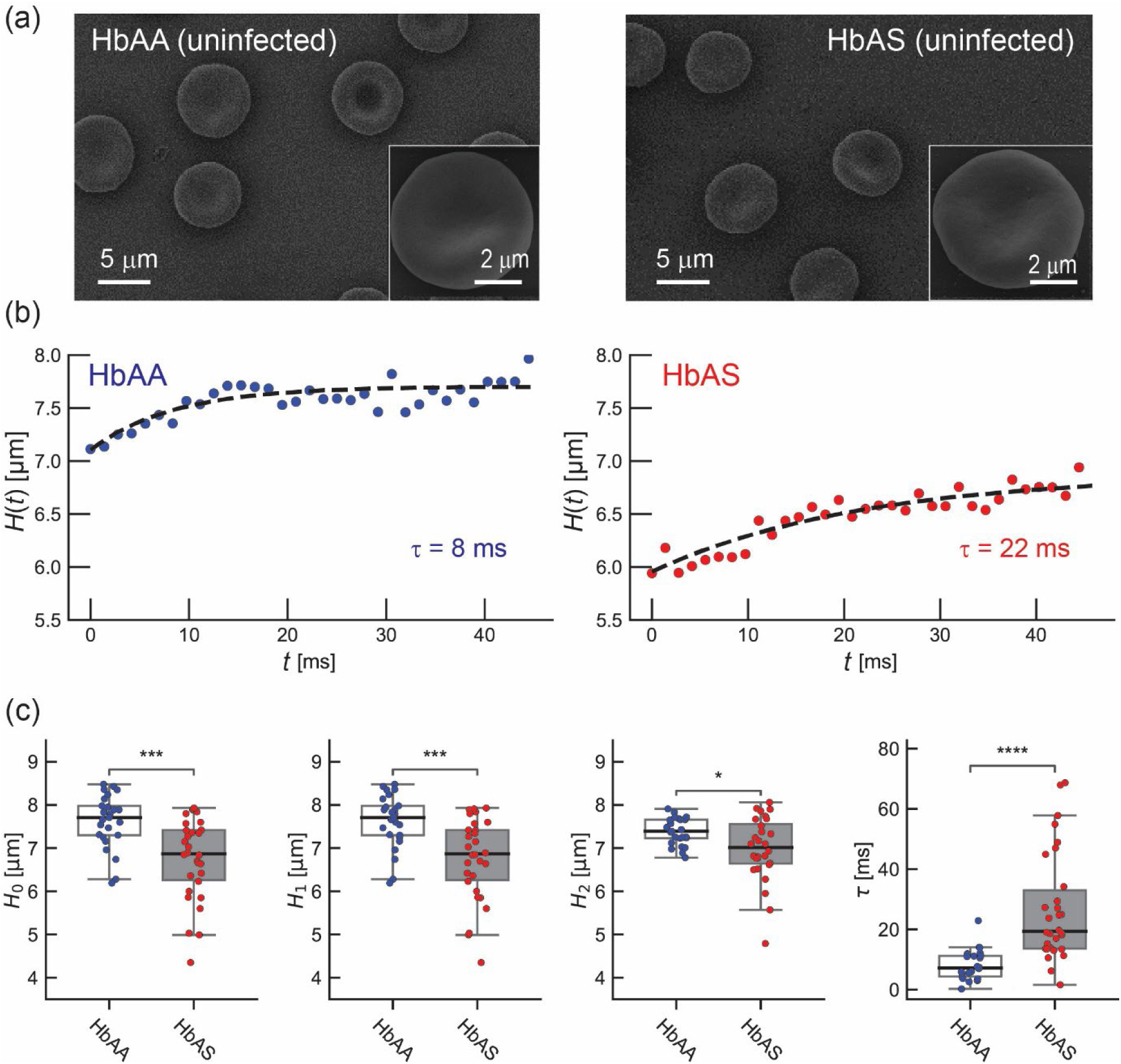
Dynamic shape recovery of uninfected HbAA and HbAS erythrocytes. (a) Scanning electron micrographs of uninfected HbAA and HbAS erythrocytes, exhibiting normal discocyte shapes. (b) Dynamic shape recovery monitored by plotting *H*(*t*) as a function of *t*. Blue; HbAA, red; HbAS. (c) Statistical comparison of *H*_0_, *H*_1_, *H*_2_, and *τ* between HbAA (*N* = 26) and HbAS (*N* = 36).

Figure 3b shows representative shape recovery curves for both cell types. Here, *t* = 0 corresponds to *t*_1_ in Figure 2c, marking the time point of complete exit from the constriction and the starting point for exponential fitting. From these curves, we extracted the parameters *H*_0_, *H*_1_, *H*_2_, and *τ* (number of cells analyzed: HbAA, *N* = 26; and HbAS, *N* = 36). The average initial cell width *H*_0_ was significantly smaller in HbAS erythrocytes compared with HbAA cells (*p* < 0.001; Student’s t-test;). This observation is consistent with previous reports showing that HbS containing erythrocytes exhibit an increased propensity for vesiculation, attributed to oxidative damage and the shedding of affected membrane fragments (Waldecker et al., 2017, Chaves et al., 2008, Hebbel, 1991). The resulting reduction in membrane area is also reflected in the minimum width at the complete exit, *H*_1_, which follows the same trend.

Notably, the characteristic recovery time *τ* differed significantly between the two cell types (**** *p* < 0.0001; Student’s t-test). The median relaxation time of HbAS erythrocytes (*τ* ≈ 20 ms) was approximately 2.5-fold higher than that of HbAA cells (*τ* ≈ 8 ms). This marked delay in shape recovery indicates altered viscoelastic properties in HbAS erythrocytes, arising from increase in membrane tension and impaired membrane–cytoskeleton coupling (Hebbel, 1991, Cyrklaff et al., 2011, Fröhlich et al., 2019). Together, these results validate our assay as a sensitive and quantitative approach to resolve subtle differences in erythrocyte deformability and recovery dynamics.

We further noted that some uninfected erythrocytes adopted a parachute-like shape upon exiting the constriction (Figure S3). This observation is in good agreement with the phase diagram calculated by computer simulations predicting that erythrocytes flowing in microchannels should exhibit a parachute-like morphology when the capillary number exceeds a critical threshold level, *C*_k_* ≈ 10^2^ (Lázaro et al., 2014). *C*_k_ is defined as the ratio of elastic to viscous relaxation times (Lázaro et al., 2014), *C_k_* = *τ*_ela_⁄*τ*_vis_, where the elastic relaxation time is given by *τ*_ela_ = *ηd*^3^⁄*κ* and the viscous timescale by *τ*_vis_ = 〈*v*〉⁄*Φ*.

To assess whether similar conditions apply in our microfluidic constriction system, we determined the average flow velocity of uninfected erythrocytes in the relaxation zone from high-speed images as 〈*v*〉 = (2.8 ± 1.7) × 10^−3^ m s^−1^. Using literature values for the bending rigidity and cytoplasmic viscosity of uninfected erythrocyte, *κ* = (2.6 ± 0.7) × 10^−19^ J and *η* = (1.2 ± 0.7) × 10^−2^ N m^2^ s (Fröhlich et al., 2019), respectively, we obtained a capillary number that exceeds *C*_k_* by approximately one order of magnitude.

To further validate this interpretation under conditions matching our experimental set-up, we performed computer simulations that explicitly account for the constriction dimensions and flow field. **Figure 4a** shows simulations incorporating SDPD hydrodynamics, which successfully recapitulated the experimentally observed parachute-like shape upon release. These results are consistent with both theoretical predictions and the estimated capillary number regime, in which viscous stresses dominate over membrane elastic restoring forces and thereby govern cell shape.

**Figure 4.**
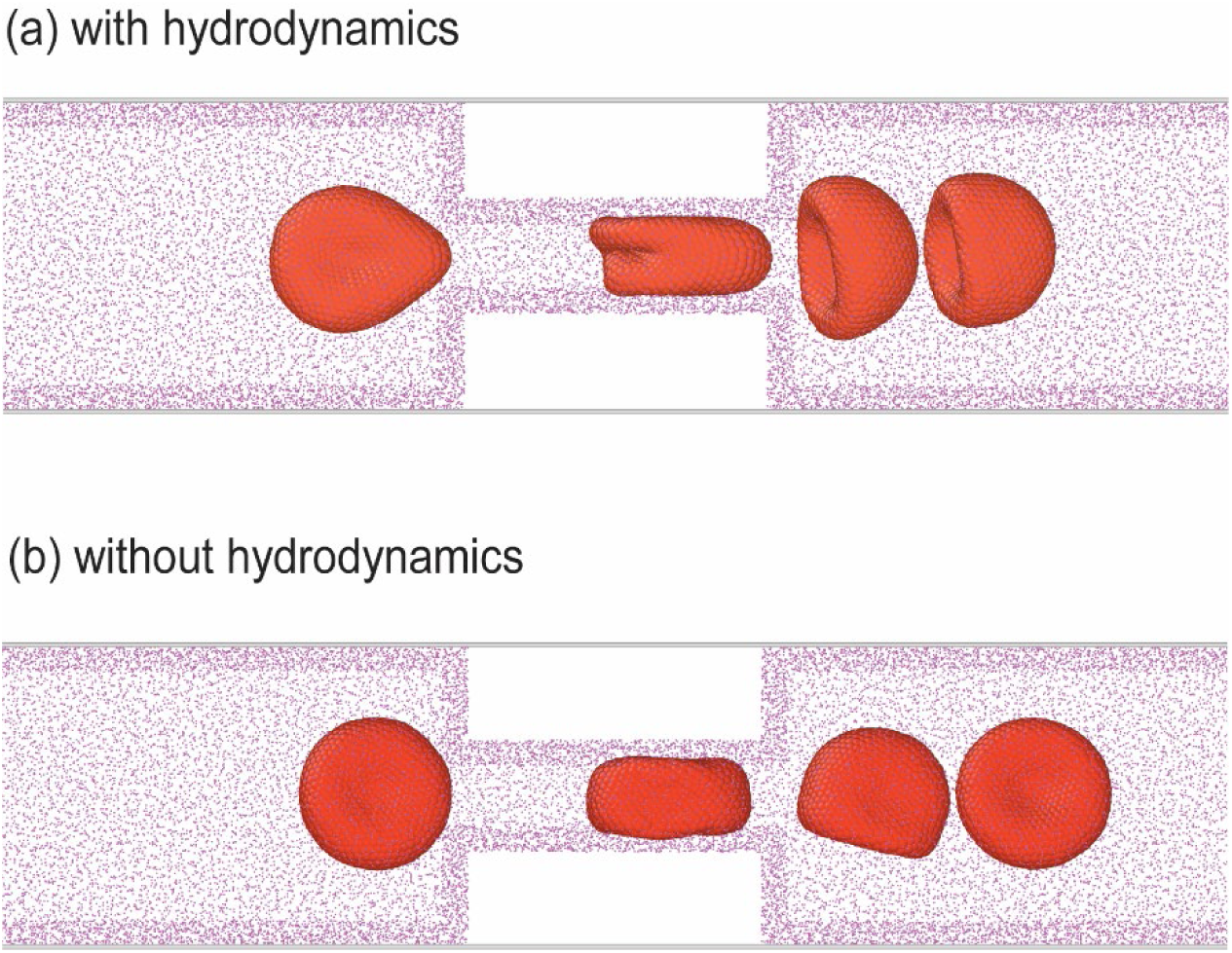
Computer simulations of viscoelastic shape recovery. Computer simulations of cells based on triangular elastic lattice networks (a) with and (b) without hydrodynamics (see more details in the main text). Experimentally observed “parachute-like” shape could be recapitulated only in the presence of SDPD hydrodynamics.

In contrast, the computer simulation that does not include hydrodynamics failed to reproduce these features and do not capture either the pronounced deformation within the constriction or the characteristic post-constriction morphology (**Figure 4b**). This discrepancy underscores the importance of fluid–structure interactions. As the cell passes through the constriction, viscous stresses and lubrication forces between the membrane and channel walls generate significant hydrodynamic friction, which contributes to both shape deformation and energy dissipation (Preira et al., 2013). Thus, accurate modeling of erythrocyte dynamics in confined flow requires explicit inclusion of hydrodynamics.

### Parasite maturation modulates shape recovery dynamics of *P. falciparum* infected wild-type HbAA erythrocytes

We next investigated HbAA erythrocytes infected with the *P. falciparum* FCR3 line at the ring, trophozoite and early schizont stage (**Figure 5a**). Infected cells at the ring, trophozoite and early schizont stages were able to successfully traverse the channels, exhibiting characteristic deformation and relaxation. In contrast, late schizont-stage infected erythrocytes consistently failed to pass and frequently became lodged within the constrictions, leading to channel occlusion. This stage-specific behavior is consistent with the progressive stiffening of cells and loss of discoidal shape during parasite maturation, resulting from both parasite growth and extensive cytoskeletal remodeling in the host (Waldecker et al., 2017, Fröhlich et al., 2019). It should further be noted that from 18 hours post invasion onwards infected erythrocytes display increasing numbers of knobs on the surface (**Figure 5b**), which contribute to membrane stiffening (Fröhlich et al., 2019, Lai et al., 2015, Quadt et al., 2012).

**Figure 5.**
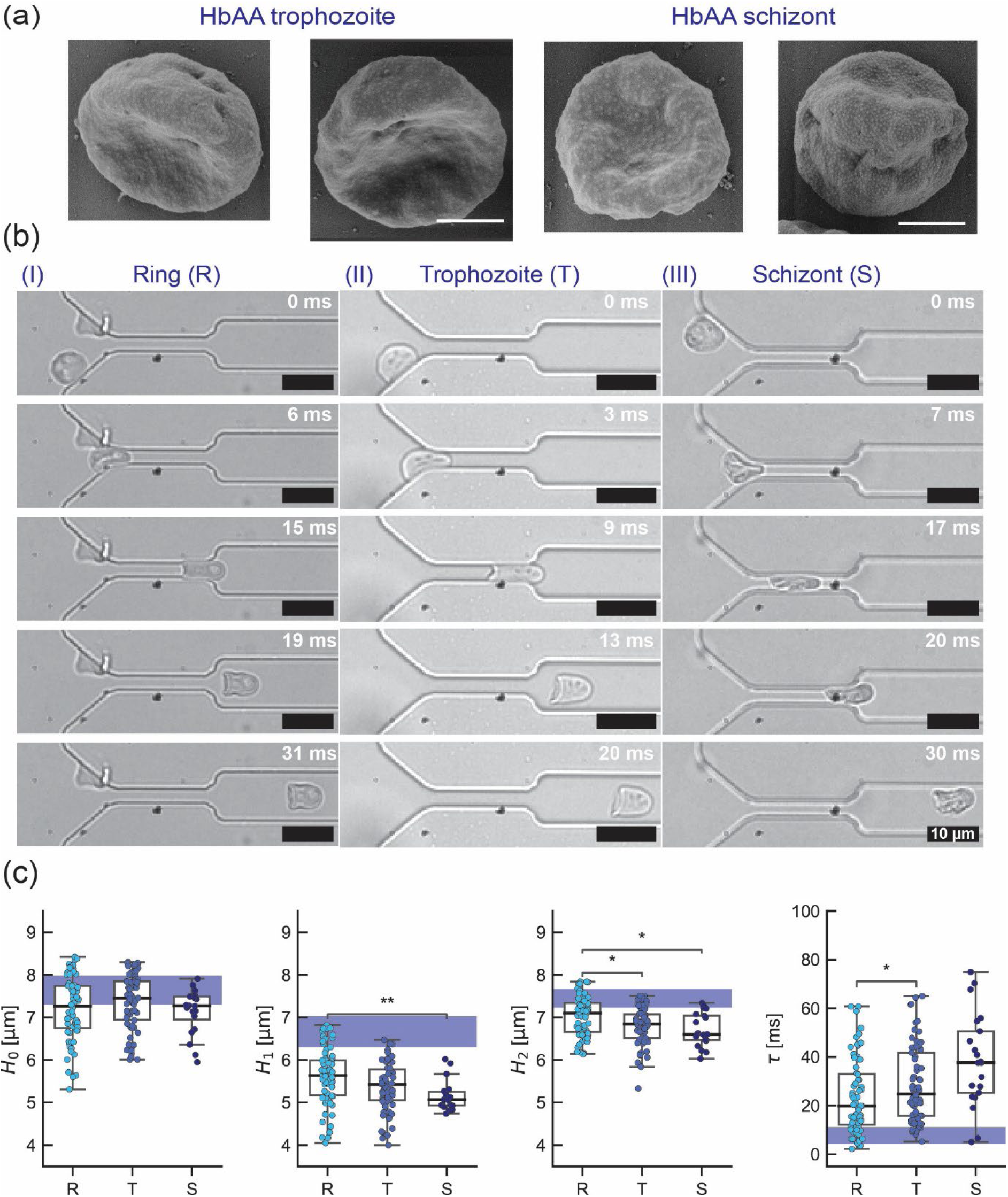
Dynamic shape recovery of *P. falciparum*-infected HbAA erythrocytes. (a) SEM images of infected HbAA erythrocytes at trophozoite state and schizont stage, showing small and uniform knobs on the surface. (b) Snapshot images of infected HbAA erythrocytes at ring, trophozoite, and schizont stages. (c) Statistical comparison of *H*_0_, *H*_1_, *H*_2_, and *τ* of HbAA erythrocytes at ring (*N* = 84), trophozoite (*N* = 77), and schizont (*N* = 22) stages. The corresponding uninfected HbAA data are highlighted in blue for visual comparison.

In the case of early schizont stage, we occasionally observed a phenomenon known as pitting, whereby the intracellular parasite is expelled while the host cell membrane remains intact (Figure S4) (Henry et al., 2020). This process was typically associated with transient deformation and localized membrane rupture followed by resealing, suggesting sufficient membrane plasticity despite ongoing remodeling. Pitting is thought to be a physiologically relevant process that occurs in the spleen, where mechanical filtration through interendothelial slits facilitates parasite removal while allowing resealed erythrocytes to return to circulation (Henry et al., 2020).

Upon passing the constriction, 80% of the ring- and trophozoite-stage infected erythrocytes (*N* = 84 and *N* = 77, respectively), exhibited a discoidal shape, whereas the remaining 20% adopted a parachute-like phenotype. In contrast, the vast majority of schizont-stage cells (>95%; *N* = 22) exited the constriction in highly deformed pancake- or bullet-like morphologies (Figure S3).

A comparative analysis of the shape recovery parameters revealed that the initial, non-perturbed cell width, *H*_0_, remained largely constant across the parasite stages and did not differ significantly from that of uninfected erythrocytes (**Figure 5c**). In comparison, the cell width at complete exit, *H*_1_, decreased progressively with parasite maturation (*p* < 0.001, Kruskal Wallis ANOVA on ranks), with ring-stage cells already showing significantly lower values than uninfected erythrocytes (*p* < 0.001, Kruskal Wallis ANOVA on ranks) (Figure 5c). This observation indicates that while ring- and trophozoite-stage cells retain partial shape recovery capacity, schizont-stage erythrocytes exhibit a pronounced impairment in their ability to re-expand after deformation.

This tendency is further accentuated in the final cell width *H*_2_, which is significantly reduced in trophozoite- and schizont-stages compared with both uninfected erythrocytes and ring stages (*p* < 0.01; Kruskal Wallis ANOVA on ranks) (**Figure 5c**). Concomitantly, the fraction of cells exhibiting pancake- and bullet-type phenotypes increased, reflecting a shift toward irreversible deformations (Figure S3).

Consistent with decreasing *H*_1_ and *H*_2_ values, the relaxation time *τ* increased significantly with parasite maturation, rising from 8 ms in uninfected erythrocytes to 20 ms in rings, and further to 26 ms and 38 ms in trophozoites and schizonts, respectively (Figure 5c). This progressive slowing of shape recovery indicates increasing viscoelastic resistance and reduced membrane adaptability.

### Distinct shape recovery dynamics of infected HbAS erythrocytes

Parasites grown in HbAS erythrocytes exhibit an aberrant knob architecture, characterized by fewer but enlarged knobs compared with infected HbAA cells (Cholera et al., 2008). In our system, the FCR3 line displayed a knob density of 17.0 ± 2.0 µm^-2^ with an average height of 93.0 ± 6.0 nm when grown in HbAS erythrocytes, compared with 24.0 ± 2.0 µm^−2^ and 44.0 ± 3.0 nm, respectively, in HbAA cells (**Figure 6a**). Infected HbAS in our shape recovery assay revealed additional features that are distinct from those observed in infected wild-type cells (**Figure 6b**). First, both trophozoite- and schizont-stage HbAS cells predominantly exhibited bullet-like phenotypes upon exiting the constriction, and no parachute-like phenotypes were observed at the schizont stage (Figure S3). Second, while the initial cell width *H*_0_ remained largely unchanged across parasite stages, the initial cell height *H*_1_ in ring stage HbAS erythrocytes (ring, *N* = 95) was preserved compared to uninfected HbAS cells (**Figure 6c**).

**Figure 6.**
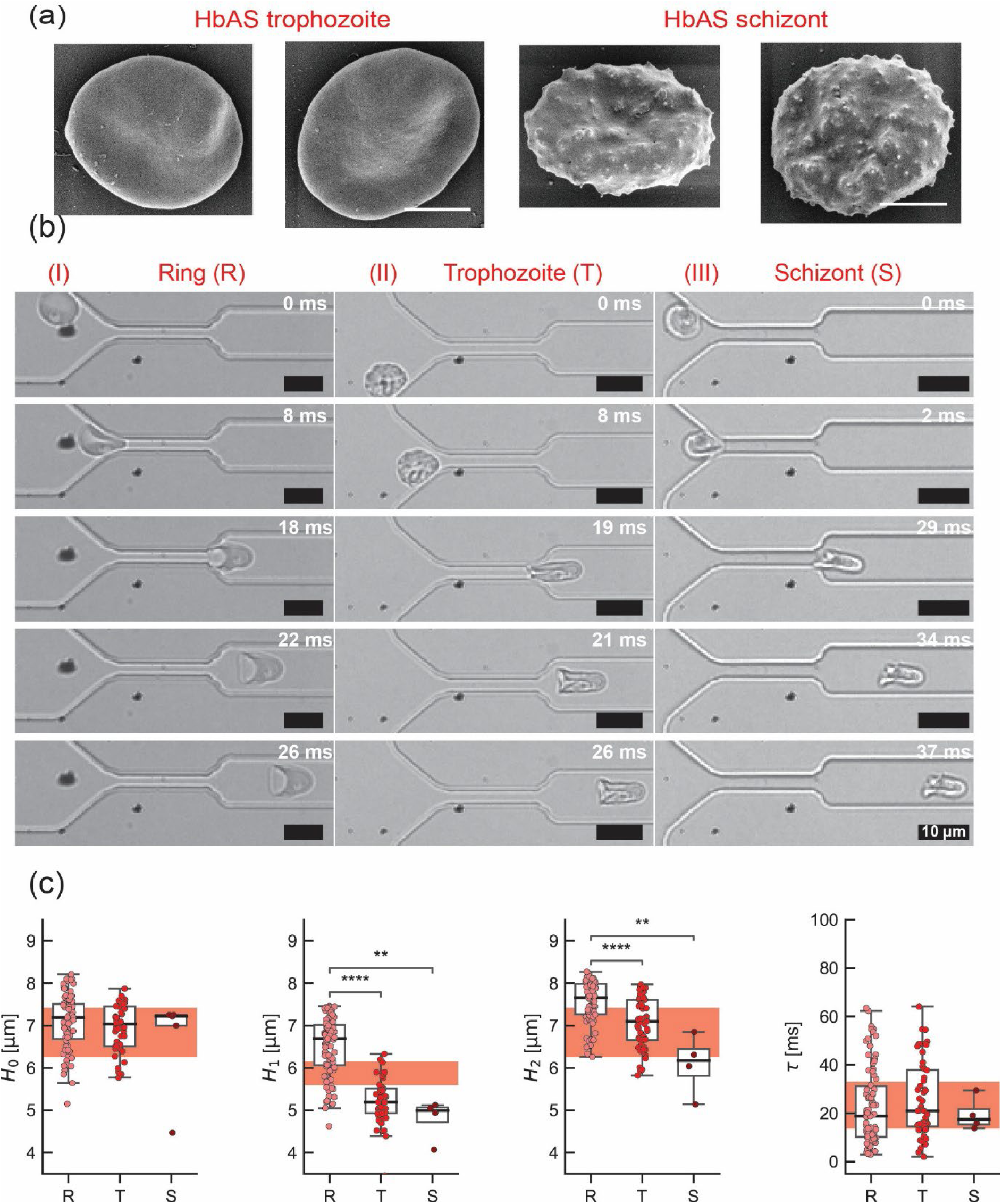
Knob phenotype and dynamic shape recovery of infected HbAS are distinct from HbAA. (a) SEM images of infected HbAS erythrocytes at trophozoite stage and schizont stage. Knobs could hardly be found on the surface of HbAS trophozoite, whereas fewer and larger knobs are harbored on the surface of HbAS schizont. (b) Snapshot images of infected HbAS erythrocytes at ring, trophozoite and schizont stages. Note that about 90 % of HbAS trophozoites showed pancake- and bullet-like phenotypes. (c) Statistical comparison of *H*_0_, *H*_1_, *H*_2_, and *τ* of HbAS erythrocytes at ring (*N* = 95), trophozoite (*N* = 63), and schizont (*N* = 5) stages. The corresponding uninfected HbAS data are highlighted in red for visual comparison.

During further parasite maturation from ring stage to trophozoite and schizont stage (trophozoite, *N* = 63; and schizont, *N* = 5), both *H*_1_ and *H*_2_ decreased progressively (**Figure 6c**), with a more pronounced reduction than in infected HbAA cells. This trend indicates that while ring stage HbAS erythrocytes preserve their shape recovery, deformation becomes increasingly irreversible in later stages of infected HbAS cells, reflecting the intrinsically altered viscoelastic properties of HbAS erythrocytes (Figure S3). This is compounded by progressive structural constraints arising from parasite growth and cytoskeletal remodeling in host cells.

Strikingly, however, the characteristic relaxation time *τ* remained approximately constant at ∼ 20 ms across all stages, showing no significant deviation from uninfected HbAS erythrocytes. This behavior contrasts with HbAA cells, where *τ* increased markedly with parasite maturation. The decoupling between increasing deformation (decreasing *H*_1_ and *H*_2_) and unchanged recovery kinetics suggests a fundamentally different modulation of viscoelastic properties in HbAS erythrocytes.

### Knob architectures modulate shape recovery dynamics

To disentangle the contribution of knob architecture from that of host cell factors—most notably hemoglobin composition—we investigated an FCR3-derived parasite mutant line, termed SR1 (**Figure 7a**, Figure S5). Figures 7b and 7c show STED and SEM images of HbAA SR1 at trophozoite stage. Figure 7d presents the snapshot images of HbAA SR1 at trophozoite stage, and *H*_0_, *H*_1_, *H*_2_, and *τ* of HbAA SR1 (*N* = 40) are statistically compared with those of parental HbAA FCR3 line (*N* = 77) in Figure 7e.

**Figure 7.**
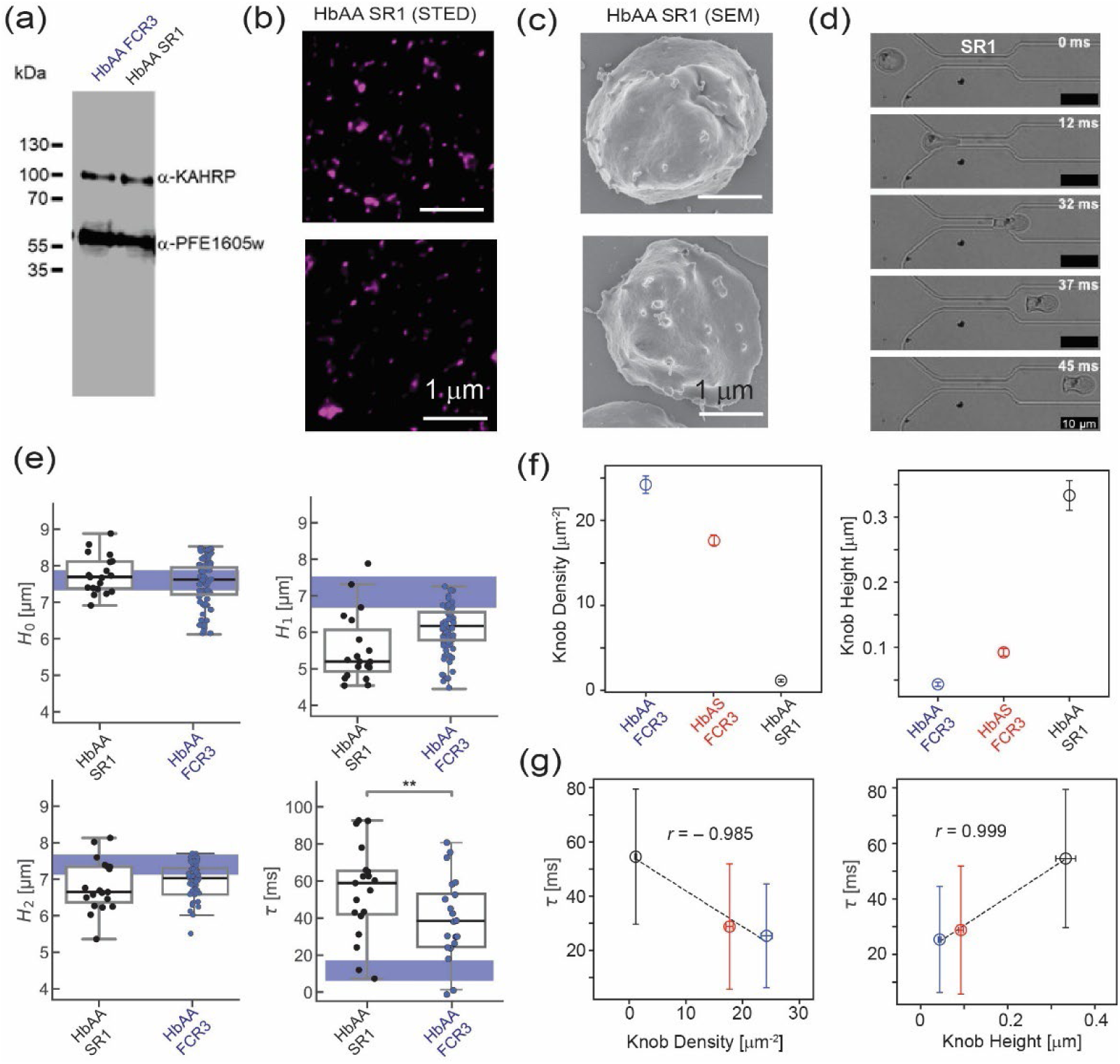
Knob architectures modulate shape recovery dynamics. (a) Western blot analysis of KAHRP expression in HbAA infected by parental FCR3 parasite and genetically modified mutant SR1 (see Figure S5 for more detail). Protein lysates were separated by SDS-PAGE and immunoblotted with primary antibody to detect KAHRP. Comparable band intensities between the parental and mutant parasites indicate that the introduced mutation does not markedly alter KAHRP protein expression. PFE1605w immunoblotted with primary antibody is also presented to demonstrate that parasite protein expression remains intact. (b) STED images of HbAA SR1 at trophozoite stage. (c) SEM images suggest about 24-fold reduction in knob density and about 7.6-fold increase in knob height compared with the parental line. (d) Snapshot images of infected HbAA SR1 erythrocytes at trophozoite stage. (e) Statistical comparison of *H*_0_, *H*_1_, *H*_2_, and *τ* of HbAA SR1 (*N* = 40) and those of parental HbAA FCR3 line (*N* = 77). The corresponding uninfected HbAA data are highlighted in blue for visual comparison. (f) Knob density (*N* = 40) and knob height (*N* = 19) of HbAA SR1 determined by SEM. For comparison, the mean and standard deviation of HBAA FCR3 (*N* ≥ 50) and HbAS FCR3 (*N* ≥ 40) are presented.

Although the SR1 mutation did not affect the geometric descriptors, *H*_0_, *H*_1_, and *H*_2_, significantly, the pronounced alterations in knob architecture had a significant impact on the relaxation time *τ*, showing almost 2-fold larger the parental line (** *p* < 0.01, Student t-test). To unravel how knob architectures modulate the shape recovery dynamics, we analyzed the SEM images and found that HbAA SR1 has about 24-fold smaller knob density (1.0 ± 1.0 knobs/µm^-2^) and 7.6-fold longer/higher knobs (333.0 ± 23.0 nm) compared with the parental HbAA FCR3 line (Figure 7f). Intriguingly, we found that the relaxation time *τ* shows a clearly negative correlation with knob density (Pearson correlation coefficient, *r* = – 0.985) and a clearly positive correlation with the knob height (*r* = + 0.999), independent from the hemoglobin composition (Figure 7g). This finding seems consistent with our previous study with knobless parasite line derived from FCR3 strain, where trophozoites showed a significantly lower membrane– cytoskeleton coupling (* *p* < 0.05) compared to the age-matched erythrocytes infected by control FCR3 (Fröhlich et al., 2019). Therefore, further studies with more mutants showing differential knob architectures will help us verify if the knob architecture, characterized by knob density and height, acts as a biomechanical regulator dominating the shape recovery dynamics of infected erythrocytes.

To place these changes in a more general context, we conducted a comparative analysis across all conditions, including uninfected erythrocytes. This analysis showed that the exit width *H_1_* decreases substantially upon infection with *P. falciparum*, with an additional reduction observed in cells infected with the SR1 mutant. The magnitude of the decrease in *H*_1_ upon infection was comparable between FCR3 parasites in trophozoite stage grown in HbAA and HbAS erythrocytes, indicating that this effect is largely independent of hemoglobin composition. In contrast, changes in *H*_0_ and *H_2_* were comparatively modest across conditions.

The analysis further highlights a pronounced increase in *τ* upon infection and underscores the influence of knob architecture on recovery dynamics. Especially, the clearly negative correlations between *τ* and knob density as well as the clearly positive correlation between *τ* and knob height independent of hemoglobin composition demonstrate that alterations in knob architecture can significantly modulate the global viscoelastic response of infected erythrocytes under large deformation conditions, such as passage through micro-vasculatures and splenic slits.

## Discussion

Our data identify *P. falciparum* infection and parasite-induced remodeling as the primary determinants of whole erythrocyte viscoelasticity and, consequently, shape recovery dynamics upon passage through a constriction. Knob architecture strongly contributes to this mechanical phenotype, while hemoglobin composition exerts an additional, though comparatively smaller, effect. This is consistent with previous reports showing that *P. falciparum*-infected erythrocytes containing hemoglobin S or the related hemoglobin C exhibit significantly larger bending rigidity and membrane–cytoskeleton coupling compared to those of erythrocytes carrying wildtype hemoglobin (Fröhlich et al., 2019, Hebbel, 1991). While this effect is clearly detectable in our assay, it is modest relative to the contributions arising from infection and parasite-driven structural remodeling. Although our setup can only address single cell behavior, these findings suggest that the induction of knobs will have a strong detrimental effect for oxygen delivery in microcirculatory flow.

Our assay probes the shape recovery of the whole cell after transient, large deformation. The extracted dynamic parameters provide direct insight into erythrocyte mechanics: the relaxation time *τ* reflects the timescale over which the viscoelastic response is governed by the interplay between cytosolic viscosity, membrane tension, bending rigidity, and membrane–cytoskeleton coupling. In parallel, the geometric parameters *H*_0_, *H*_1_, and *H*_2_ report on the significance of deformation and the reversibility of shape recovery. Together, these observables define a quantitative framework for assessing erythrocyte mechanics beyond equilibrium or small-deformation regimes typically accessed by conventional techniques.

Uninfected HbAS erythrocytes exhibited a markedly slower shape recovery compared with HbAA cells, earmarked by the increased relaxation time. This behavior is consistent with a baseline alteration of membrane mechanics associated with the presence of hemoglobin S (Hebbel, 1991). Oxidative processes linked to HbS auto-oxidation promote modifications of key membrane components, including band 3 clustering and spectrin crosslinking (Ferru et al., 2011, Gwozdzinski et al., 2026), which impair membrane–cytoskeleton coupling and increase the effective shear modulus of the cell membrane (Fröhlich et al., 2019). In addition, increased cytosolic viscosity and membrane loss through vesiculation further contribute to the altered viscoelastic response (Hebbel, 1991). These findings indicate that hemoglobin composition establishes a pre-conditioned mechanical state that persists independently of infection.

Upon infection, erythrocytes undergo extensive remodeling driven by parasite growth and protein export into the host cell compartment (Jonsdottir et al., 2021, Maier et al., 2009, Wiser, 2023). In HbAA cells, this remodeling is reflected in a progressive increase in relaxation time from ring to trophozoite to schizont stages, consistent with increasing membrane tension and enhanced spectrin–membrane coupling. This progressive increase in *τ* can be explained by gradual stiffening of the composite membrane, driven by parasite-induced processes such as formation and maturation of protein knobs, as well as reorganization of the host cytoskeleton (Cyrklaff et al., 2011, Fröhlich et al., 2019, Wiser, 2023).

In contrast, infected HbAS erythrocytes exhibited a distinct mechanical response. In ring stage, the deformation is not significantly altered by the parasite when compared to uninfected HbAS erythrocytes. In HbAA erythrocytes progressive stiffening during ring-stage infection can primarily be attributed to the ring-infected erythrocyte surface antigen (RESA) protein, which are released by merozoites and becomes associated with the erythrocyte membrane at the time of invasion (Diez-Silva et al., 2012, Mills et al., 2007). RESA is phosphorylated by a host associated erythrocyte membrane kinase (Foley et al., 1990) and binds to the spectrin tetramer increasing its stability (Pei et al., 2007). Infected HbAS erythrocytes show an altered phosphorylation pattern compared to infected HbAA erythrocytes which might influence the ability of RESA to interact with its spectrin binding side (Chauvet et al., 2021). This altered interaction could account for the preserved deformability observed in ring stage HbAS erythrocytes.

While the fraction of irreversible deformation increased markedly with further parasite maturation, as indicated by the decrease in *H*_1_ and *H*_2_, the relaxation time remained largely unchanged. In light of a recent optical tweezer study (Gironella-Torrent et al., 2024) that indicated that fast (0.01-0.1s), intermediate (4s) and slow (70s) relaxation processes are related mainly to membrane, cytoskeleton and cytoplasm, respectively, our results suggest that upon exit from a constriction, mainly the two first elements contribute, while the effect of the cytoplasm should not be relevant on the short time scale of tens of ms. This explains why the infection, which strongly changes the coupling between membrane and cytoskeleton, leads to a clear increase in relaxation times, while the HbAS cells, which suppress these changes but have altered solution characteristics, do not show a strong change in relaxation times.

A key structural correlate of this behavior is the altered knob architecture in HbAS erythrocytes. At the trophozoite stage, HbAS cells display a substantially reduced knob density compared with HbAA cells, while the remaining knobs are larger and more irregularly distributed (Cholera et al., 2008, Sanchez et al., 2019, Fröhlich et al., 2019, Fairhurst et al., 2012). Electron tomography studies further indicate that this altered organization leads to weakened and spatially heterogeneous coupling between knobs and the underlying spectrin network (Cyrklaff et al., 2011, Cyrklaff et al., 2016, Fröhlich et al., 2019). As a consequence, the mechanical reinforcement typically associated with knob formation in HbAA erythrocytes is diminished or redistributed in HbAS cells, resulting in a different balance between viscous and elastic responses.

Consistent with this interpretation, flicker spectroscopy studies have shown that the evolution of mechanical parameters during parasite maturation differs substantially between HbAA and HbAS erythrocytes (Fröhlich et al., 2019). In HbAS cells, the increase in bending modulus from the uninfected to the trophozoite stage is more pronounced, likely reflecting the accumulation of membrane-bound oxidized hemichromes and oxidative modifications of membrane skeletal components (Gwozdzinski et al., 2026, Hebbel, 1991, Cyrklaff et al., 2011). In contrast, changes in membrane tension and spectrin–membrane coupling follow distinct trajectories compared with HbAA and other hemoglobinopathies such as HbAC, highlighting pronounced increases in membrane tension (Fröhlich et al., 2019). Computational modeling further supports the notion that both knob density and spectrin–membrane coupling critically influence the effective mechanical response of infected erythrocytes.

The contribution of knob architecture was further examined using the SR1 mutant line, which exhibits enlarged and sparsely distributed knobs in HbAA erythrocytes. These pronounced alterations in knob morphology, compared with the parental FCR3 line, significantly affected shape recovery dynamics, as reflected by a prolonged relaxation time *τ*. These findings demonstrate that knob architecture can modulate the global viscoelastic response of the cell under large-deformation conditions.

Mechanistically, enlarged knobs are likely to introduce localized stiffening and increase resistance to membrane re-expansion, while reduced knob density may lead to a more heterogeneous distribution of membrane–cytoskeleton anchoring points. This combination is expected to slow down relaxation dynamics. Notably, the effects on geometric parameters (*H*_0_, *H_1_*, and *H*_2_) are comparatively modest relative to the pronounced increase in *τ*, suggesting that knob architecture primarily influences recovery kinetics rather than the extent of deformation. This distinction suggests that knobs modulate how mechanical stresses are dissipated and redistributed across the membrane–cytoskeleton composite, rather than solely determining static deformability.

These findings have important implications for understanding the protective effect of the sickle cell trait against severe malaria (Cholera et al., 2008). Altered mechanical properties of HbAS erythrocytes, both at baseline and upon infection, are likely to influence their behavior in the microcirculation. Increased irreversibility of deformation and altered recovery dynamics may promote splenic clearance of infected cells, while aberrant membrane remodeling and reduced cytoadhesive capacity may impair sequestration. Together, these effects could contribute to reduced parasite survival and disease severity in HbAS individuals.

Several limitations should be considered. The use of an ellipse-based representation simplifies complex cell shapes and may not fully capture local deformations, particularly in highly irregular cells. In addition, while the microfluidic geometry mimics key aspects of microcirculatory constraints, it does not fully reproduce the biochemical and mechanical complexity of the *in vivo* environment, particularly within the spleen. Nevertheless, the consistency of our results across multiple conditions supports the robustness of the assay.

In summary, our study establishes dynamic shape recovery as a sensitive and quantitative readout of erythrocyte mechanics. By linking hemoglobin composition, parasite-induced remodeling, and knob architecture to distinct viscoelastic signatures, we provide new insight into the biomechanical alterations underlying malaria pathophysiology. Future work may extend this approach to other hemoglobinopathies, pharmacological perturbations, and genetically modified parasite lines, with the aim of further dissecting the interplay between host cell mechanics and parasite virulence.

## Author Contributions

M.T., U.S.S., and M.L. designed the research and supervised the project. J.C., V.L., and C.P.S. performed the experiments. J.C., V.L., P.R.H., L.L., M.L., and M.T. analyzed the data. A.K.D. performed the computer simulations with help from U.S.S.; M.L. and M.T. wrote the original manuscript. All authors participated in discussion and manuscript editing.

## Acknowledgement

J.C. thanks A. Yamamoto for inspiring discussion for the image analysis, and M.T. and J.C. thanks HeKKSaGOn Alliance for supports. This work was supported by the Deutsche Forschungsgemeinschaft through the Collaborative Research Center SFB 1129 (project number 240245660, F.H., U.S.S., M.L., and M.T.). M.T. thanks Nakatani Foundation for support.

